# Navigating the Urban River: Transcriptomic Responses of Freshwater Fish to Multiple Anthropogenic Stressors

**DOI:** 10.1101/2024.08.19.608252

**Authors:** Camilo Escobar-Sierra, Kathrin P. Lampert

**Affiliations:** Institute of Zoology, Universität zu Köln Mathematisch-Naturwissenschaftliche Fakultät, Zülpicher Str. 47b, Köln, NRW, 50674, Germany

**Keywords:** Pollution, Gene expression, Aquatic Ecosystem, Physiology, Molecular ecology

## Abstract

Urbanization imposes multiple anthropogenic stressors on freshwater ecosystems, affecting aquatic species’ physiological responses. This study explores the transcriptomic responses of the freshwater fish *Cottus rhenanus* to various stressors in an urban river system. RNA sequencing of fish from multiple stations revealed significant seasonal variations in gene expression, with a higher number of differentially expressed genes (DEGs) observed in the summer. Fish at the station experiencing the highest anthropogenic pressure showed notable responses, particularly during warmer months, with enriched pathways related to metabolism, oxidative stress, and immune responses. Key findings include the activation of metabolic stress pathways and immune system genes, such as IL-17 and MAPK pathways, influenced by high temperatures, salinity, and low oxygen levels. Pathway enrichment analyses highlight the impact of temperature and salinity on oxidative stress and osmoregulation, revealing the critical role of the transportome in adapting to salinity changes. These findings showcase the complex interactions between stressors and physiological responses, emphasizing the need for integrated conservation strategies to manage urban stream ecosystems.

## Introduction

Urbanization is set to significantly increase, with the United Nations Population Division (UNPD) projecting an expansion of urban land cover by 1.2–1.8 million km² by 2030 (UNPD, 2019). This rapid growth has severe implications for freshwater habitats and biodiversity, a phenomenon often described as “Urban Stream Syndrome.” This syndrome encapsulates various impacts on streams, including altered channel morphology, highly variable hydrographs, reduced biotic richness, and elevated levels of nutrients and contaminants (McDonald et al., 2019; C. J. Walsh et al., 2005). Additionally, urban streams suffer from nonpoint source pollution and climate variability, leading to complex “chemical cocktails” that act as multiple stressors (Kaushal et al., 2019; Schäfer et al., 2023). These multiple stressors pose significant threats to freshwater life, biodiversity, and ecosystem stability (Johnson & Penaluna, 2019), contributing to riverine biodiversity loss and altered community structures (Barrett et al., 2022). Environmental fluctuations in water quality, driven by anthropogenic stressors such as pH, salinity, temperature, dissolved oxygen (DO), and pollution, are recognized as major drivers of the distribution and health of fish species (Bernhardt et al., 2020; Menon et al., 2023).

The Ruhrgebiet region in Western Germany where the Emscher river flows, serves as a prime example of multiple stressors affecting urban streams. As one of Europe’s most densely populated regions (Moos et al., 2021), it exhibits many factors associated with urban stream syndrome, such as channelization, reduced lateral and longitudinal connectivity, and poor water quality. Decades of wastewater discharge, coal mining, and heavy industrial activity have led to severe hydromorphological changes and pollution, with streams historically used as open sewers (Gerner et al., 2018). Coal mining has been one of the main stressors, significantly contributing to water quality issues, increasing chloride concentrations to levels as high as 3500 mg/L, far exceeding typical natural levels of less than 20 mg/L (Hintz & Relyea, 2019; Petruck & Stöffler, 2011). Although management measures and mine closures have reduced these concentrations to below 400 mg/L (Petruck & Stöffler, 2011), abandoned mines continue to contribute to freshwater salinization and chemical contamination (Schulz & Cañedo-Argüelles, 2019). To address this prolonged degradation, a large-scale restoration project initiated in 1992 aimed to improve water quality and reshape river morphology (Winking et al., 2016). As a result, some Emscher tributaries, such as the Boye, have been sewage-free since 2017, and significant improvements in channel morphology across the network have been achieved over the past 20 years (Gillmann et al., 2023). Chloride concentrations have also decreased in the Boye, although they can still be high reaching 26–286 mg/L, along with electrical conductivities between 0.4–1.93 mS cm^−1^ (Madge Pimentel et al., 2024). Despite these efforts, not all of the Emscher catchment has been restored, and main sections near Dortmund are still heavily influenced by urbanization, with high potential for increased runoff from road salts and sewage, further impacting water quality (Dugan et al., 2017).

Water quality is a critical factor for fish distribution and habitat suitability, influenced by variables such as temperature, conductivity, dissolved oxygen, and salinity. Conductivity, which correlates with the concentrations of ions like HCO3−, SO42−, and Cl−, tends to be elevated in disturbed catchments (Kefford et al., 2023). In urbanized environments, hypoxia (a significant global water pollution issue) exacerbates sublethal effects such as endocrine disruption and oxidative stress (Abdel-Tawwab et al., 2019; Pollock et al., 2007; Zhu et al., 2013). Fish, as ectotherms, are particularly vulnerable to temperature variability. Chronic temperature increases can compromise their ability to cope with additional stressors (Alfonso et al., 2021). Rapid temperature increases can induce acute stress responses, potentially impacting ecological dynamics. Freshwater salinization poses a significant threat by increasing stress or mortality among freshwater organisms, thereby affecting biodiversity and ecosystem functionality (Cunillera-Montcusí et al., 2022). Freshwater fish, adapted to low ion concentrations, must expend considerable energy on osmoregulation, with increased salinity raising this energy expenditure (Guh et al., 2015; Tseng & Hwang, 2008). Elevated salinity impacts development, growth, and respiration, with fish often avoiding high salinity levels. Fish gills, in constant contact with the water, play crucial roles in oxygen uptake, osmotic regulation, temperature regulation, and immune response, making them sensitive indicators of physiological stress (Evans et al., 1999; Jeffries et al., 2021). The co-occurrence of multiple stressors in urban rivers makes it challenging to assess of their individual and combined effects (Orr et al., 2024). Furthermore, the complex interactions of multiple stressors complicate evaluating their effects on wild fish physiological responses under real-life conditions.

Traditional methods for evaluating the effects of multiple stressors, such as endpoint mortality, laboratory manipulations, and population studies, have limitations. High-throughput methods like transcriptomics offer a comprehensive approach to assessing gene expression in organisms exposed to multiple stressors (Lowe et al., 2017). Transcriptomics can provide detailed insights into the physiological status of species by measuring responses to multiple stressors (Jeffries et al., 2021). This approach has been successfully used in previous research to identify the stressors affecting wild fish populations (Escobar-Sierra et al., 2024; Escobar-Sierra & Lampert, 2024; Jeffrey et al., 2023; Komoroske et al., 2016). Gene expression data obtained under natural conditions can reveal the molecular pathways affected by stressors, helping to identify the specific stressors influencing the physiological response of fish in multiple stressor scenarios.

A species that is sensitive to multiple stressors is *Cottus rhenanus* (Markert et al., 2024a). In the Emscher catchment it was found in relicts, less affected river stretches but has been successfully reintroduced into several sites (Stemmer & Jacobs, 2015). *Cottus* generally require low water temperatures and high oxygen levels, restricting them to smaller, well-oxygenated cold streams (Brown, 1989; S. J. Walsh et al., 1997). Their sensitivity and conservation status in the Emscher catchment make them an ideal species to assess the effects of multiple stressors in urban streams.

The primary objective of this study, therefore, is to assess the physiological response of *Cottus rhenanus* to multiple anthropogenic stressors across the Emscher river catchment during both warm and cold seasons. This research aims to elucidate the relationship between these stressors and the gene expression profiles of gill tissue, identifying enriched physiological pathways and their interconnections. It is hypothesized that *Cottus rhenanus* in areas with higher anthropogenic stress will exhibit distinct gene expression profiles, enriched in pathways related to stress response, immune function, growth, and reproduction, regardless of the season. High-throughput mRNA sequencing and differential gene expression analyses will be used to examine these profiles along a gradient of stressors, with multivariate statistical methods exploring the relationships between gene expression and enriched pathways. This study employs de novo transcriptomic analysis to provide insights into the molecular mechanisms driving the physiological responses of a non-model wild fish species to multiple stressors in an urban stream environment.

Conducted under field conditions where multiple stressors interact, this research emphasize the importance of measuring physiological responses in wild, non-model organisms. By integrating transcriptomic insights into broader ecological assessments and management practices, this study highlights the potential of transcriptomic approaches in evaluating responses to both known and unknown stressors. This approach demonstrates their value in assessing the overall status of wild fish and guiding conservation strategies. Ultimately, this research supports the assertion that transcriptomics is essential for providing insights into complex stressor interactions and enabling informed conservation strategies. It enhances our understanding of how freshwater species adapt to complex environmental challenges, and it paves the way for more accurate assessments of physiological status in organisms, with significant implications for conservation. Future research should focus on refining rapid screening tools using candidate genes and exploring the physiological responses to multiple stressors across various species and environments, thereby improving the management of endangered species under multiple anthropogenic stressors.

## Material and methods

### Sampling design

Six stations in wadeable sections of the Emscher river catchment were sampled for *Cottus rhenanus* in the warm (June) and cold periods (October) of the year 2022 (Figure 1) and a set of water chemistry, and hydro-geomorphological variables were taken. Water temperature, pH, conductivity, and dissolved oxygen (DO) were measured in the field using a multiparametric probe and shading, straightening and substrate diversity (SDV) were determined following the water framework directives and expert decision. Warter level at pole is the water depth in the mid-section of the transversal profile of stream. Water samples were taken in dark tinted bottles and frozen under -20 °C and taken to the lab where the concentrations of nitrate, chloride and sulfate were measured. The table with values of the recorded variables is presented in Table 1.

**Figure 1.**
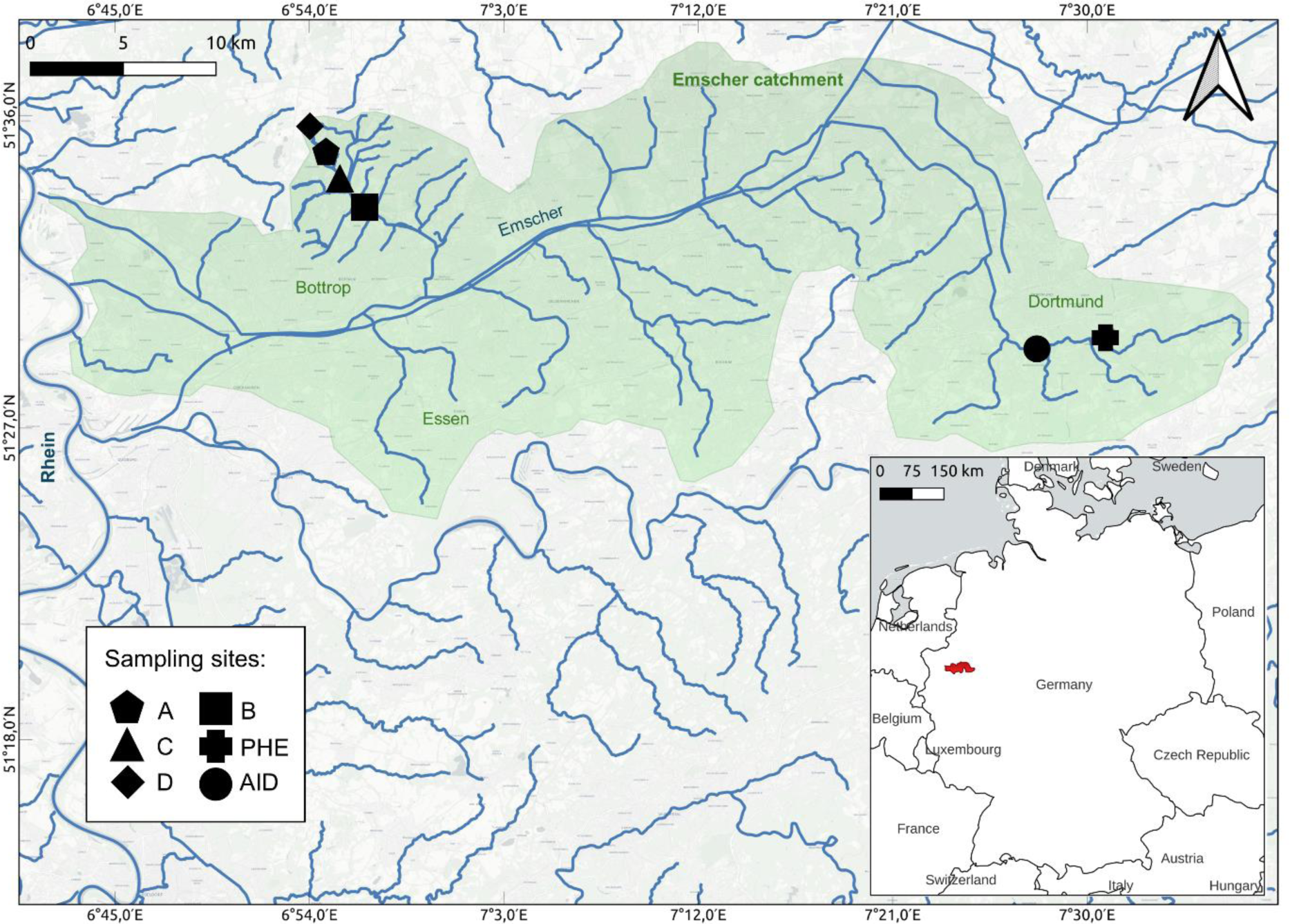
Fish sampling stations in the Emscher river catchment. Three fish were collected from each site for gill tissue RNAseq analysis. Sampling occurred once in the warm season (June) and once in the cold season (October).

**Table 1.**
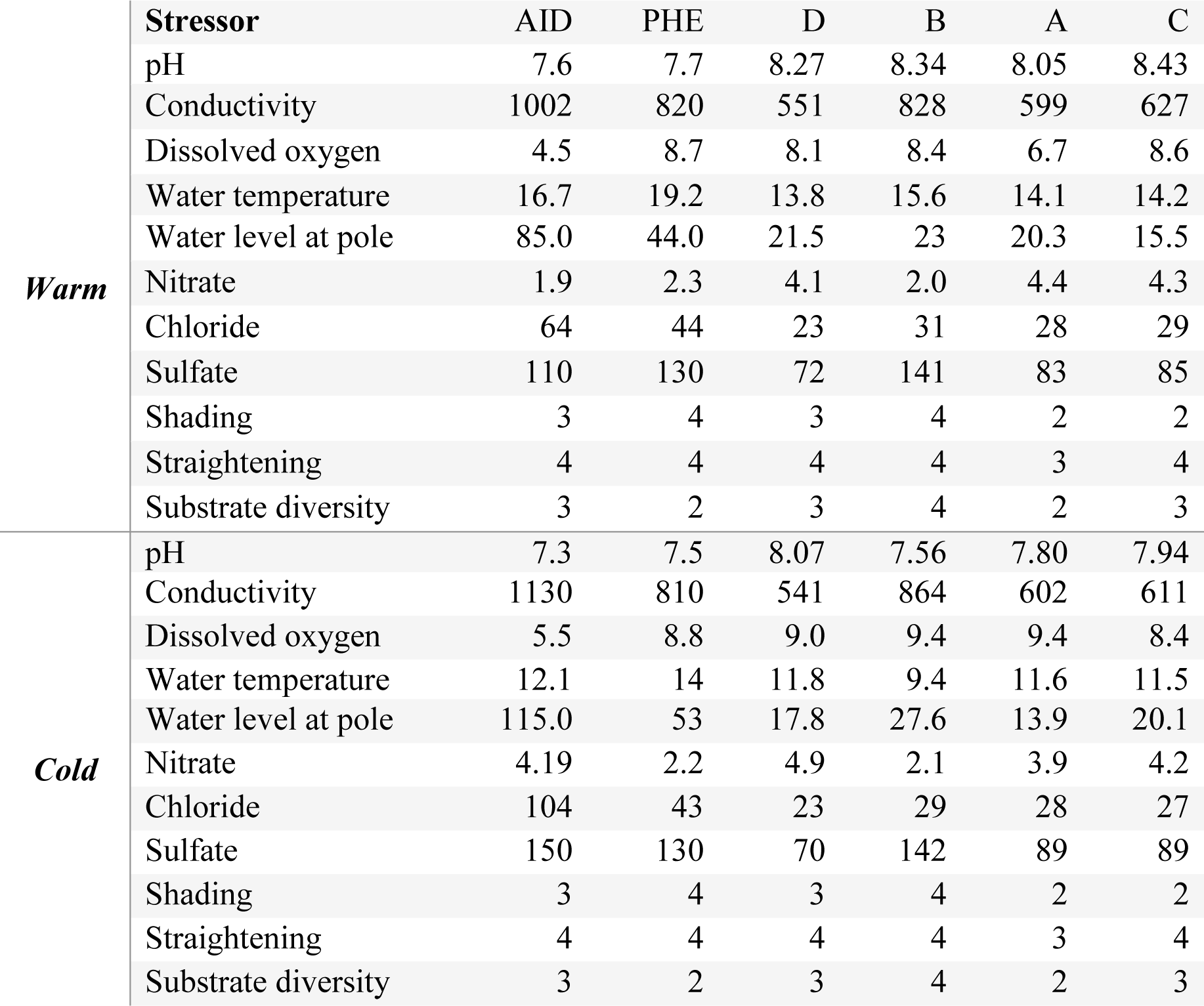
Summary of the stressor measured in all station in both seasons of sampling. Shading, straightening and Substrate diversity values range from 1 (best) to 5 (worst). Water level at pole measured in cm. Conductivity in µS/cm. Dissolved oxygen, nitrate, chloride, sulfate in mg/l.

|  | **Stressor** | **AID** | **PHE** | **D** | **B** | **A** | **C** |
| --- | --- | --- | --- | --- | --- | --- | --- |
| **Warm** | pH | 7.6 | 7.7 | 8.27 | 8.34 | 8.05 | 8.43 |
|  | Conductivity | 1002 | 820 | 551 | 828 | 599 | 627 |
|  | Dissolved oxygen | 4.5 | 8.7 | 8.1 | 8.4 | 6.7 | 8.6 |
|  | Water temperature | 16.7 | 19.2 | 13.8 | 15.6 | 14.1 | 14.2 |
|  | Water level at pole | 85.0 | 44.0 | 21.5 | 23 | 20.3 | 15.5 |
|  | Nitrate | 1.9 | 2.3 | 4.1 | 2.0 | 4.4 | 4.3 |
|  | Chloride | 64 | 44 | 23 | 31 | 28 | 29 |
|  | Sulfate | 110 | 130 | 72 | 141 | 83 | 85 |
|  | Shading | 3 | 4 | 3 | 4 | 2 | 2 |
|  | Straightening | 4 | 4 | 4 | 4 | 3 | 4 |
|  | Substrate diversity | 3 | 2 | 3 | 4 | 2 | 3 |
| **Cold** | pH | 7.3 | 7.5 | 8.07 | 7.56 | 7.80 | 7.94 |
|  | Conductivity | 1130 | 810 | 541 | 864 | 602 | 611 |
|  | Dissolved oxygen | 5.5 | 8.8 | 9.0 | 9.4 | 9.4 | 8.4 |
|  | Water temperature | 12.1 | 14 | 11.8 | 9.4 | 11.6 | 11.5 |
|  | Water level at pole | 115.0 | 53 | 17.8 | 27.6 | 13.9 | 20.1 |
|  | Nitrate | 4.19 | 2.2 | 4.9 | 2.1 | 3.9 | 4.2 |
|  | Chloride | 104 | 43 | 23 | 29 | 28 | 27 |
|  | Sulfate | 150 | 130 | 70 | 142 | 89 | 89 |
|  | Shading | 3 | 4 | 3 | 4 | 2 | 2 |
|  | Straightening | 4 | 4 | 4 | 4 | 3 | 4 |
|  | Substrate diversity | 3 | 2 | 3 | 4 | 2 | 3 |

The stations were chosen to encompass the gradient of anthropogenic stress representative of the matrix of impacts that can be found in the Emscher catchment, from the mostly agricultural reaches in the headwaters and tributaries to some of the highly urbanized reaches in the main channel of river. Stations AID and PHE in the Emscher main channel and located in a densely urbanized area in Dortmund represent strong levels of anthropogenic stress. These stations exhibit the highest electrical conductivity, chloride, temperature, water levels, and sulfate, alongside lower dissolved oxygen, indicating significant stress in both cold and warm seasons. Station B is in the middle section of the Boye river, a tributary of the Emscher, this location has intermediate level of anthropogenic stress. Stations A, C and D are located towards the headwaters of the Boye and represent some of the best water quality and lowest level of anthropogenic stress in the Emscher catchment (Figure 2 and Figure 3).

**Figure 2.**
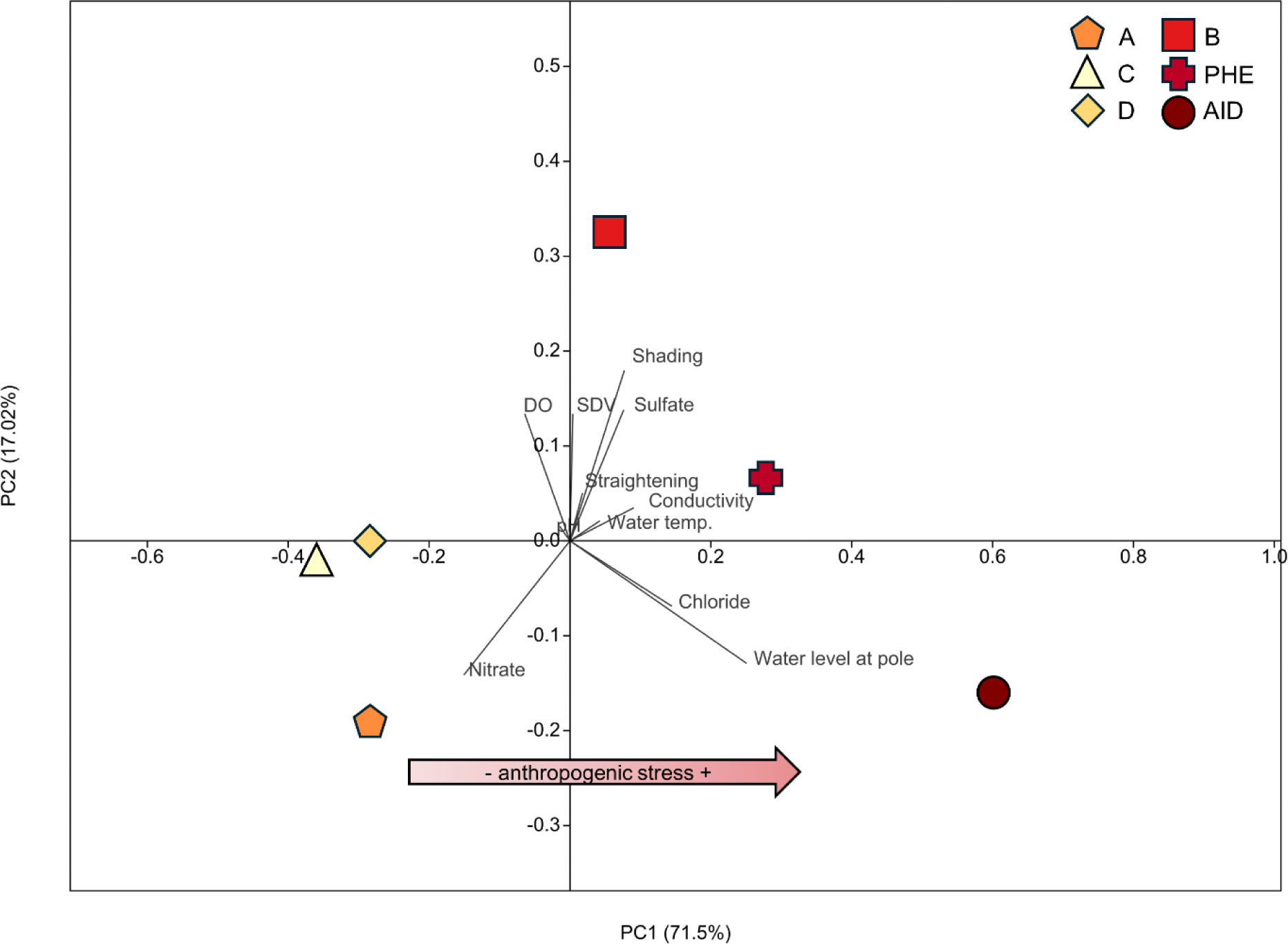
PCA of environmental variables in the Emscher river stations during the warm season. PC1 explains 71.5% of the variation and PC2 accounts for 17.02%. The bottom arrows indicate the gradient of anthropogenic stress across stations. Stations A, C, and D, characterized by higher dissolved oxygen and lower conductivity and chloride concentrations, are grouped to the left. Stations PHE and AID, with higher conductivity, chloride levels, water temperature, straightening, and lower dissolved oxygen, are grouped to the right. Station B represents mid levels of anthropogenic stress.

**Figure 3.**
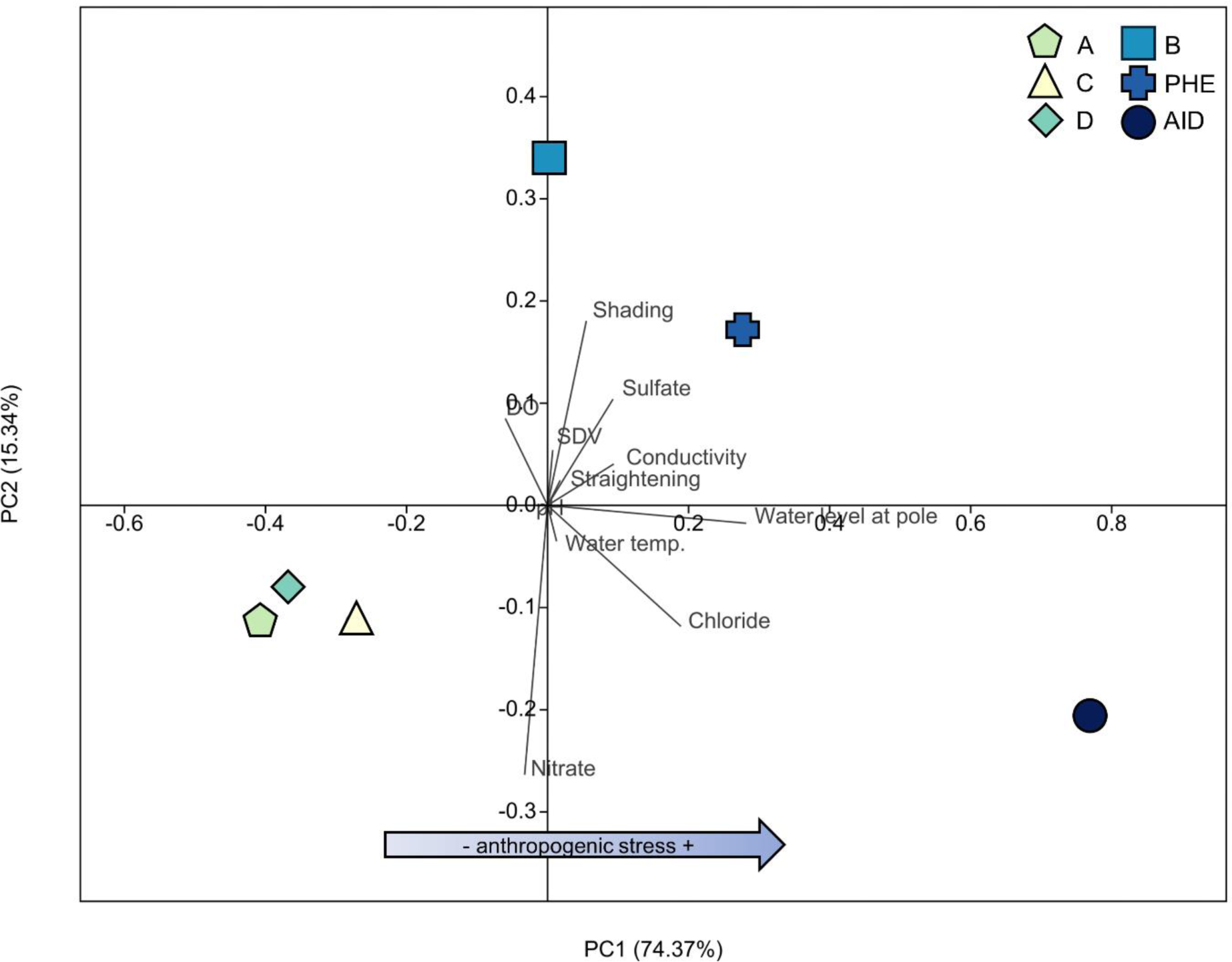
PCA of environmental variables in the Emscher River stations during the cold season. PC1 explains 74.37% of the variation and PC2 accounts for 15.34%. The bottom arrows indicate the gradient of anthropogenic stress across stations. Stations A, C, and D, characterized by higher dissolved oxygen and lower conductivity and chloride concentrations, are grouped to the left. Stations PHE and AID, with higher conductivity, chloride levels, water level, water temperature, straightening, and lower dissolved oxygen, are grouped to the right. Station B represents mid levels of anthropogenic stress.

*Cottus rhenanus* individuals were sampled using electrofishing equipment using a portable unit that generated up to 200 V and 3 A pulsed direct current (DC) in an upstream direction in all six stations in both the warm and cold seasons. All procedures were conducted following the European Directive for animal experimentation (2010/63/EU). Fishing and electrofishing permit for stations A, B, C and D in the Boye river were given by the Bezirksregierung Münster (“DE 4407-301_Kirchheller Heide”) and the sampling and permits in the Emscher main section stations (AID and PHE) were done directly by representatives of the local environmental agency (Emschergenossenschaft) and local fisheries office (Bezirksregierung Arnsberg-Obere Fischereibehörde). After capture, fish were promptly euthanized with an overdose of tricaine methanesulfonate (MS-222 at 1g./L) followed by harvesting the gill tissue, immediately fixing it in RNAprotect (QIAGEN), and storing it at -20°C. In total 36 samples were obtained, three from each station and season. The fish selected for sequencing were all males of the same size class, with similar lengths (67.77 mm ± 7.59 mm, N=36).

#### RNA Isolation and Illumina Sequencing

RNA was isolated by homogenizing fixed gill tissue samples in 700 μl RLT buffer with a 10μl:1ml concentration of β-mercaptoethanol using a FastPrep-24™ bead beater for 30 seconds at 5 m/s. The RNA was extracted with the RNeasy Mini Kit (QIAGEN), following the manufacturer’s instructions. Quality of RNA extractions was confirmed using a Nanodrop 1000 Spectrophotometer (Peqlab Biotechnologie, Germany), ensuring a concentration >50 ng/μl, an OD 260/280 ratio of 1.8-2.1, and an OD 260/230 ratio >1.5. RNA Integrity Number (RINe) values were assessed using the RNA ScreenTape system in an Agilent 2200 TapeStation, confirming RINe scores >7.0. Sequencing libraries were generated using the Illumina Tru-Seq™ RNA Sample Preparation Kit (Illumina, San Diego, CA, USA) per the manufacturer’s protocol. Illumina sequencing was performed on an Illumina HiSeq-2500 platform, generating 100 bp paired-end reads with a read depth of 30 million reads, as recommended by the Cologne Center for Genomics (CCG), Germany, based on CCG’s estimation that this read depth suffices for constructing a de novo transcriptome with 36 biological replicates.

#### Sequence Quality Control, De Novo Assembly, and Annotation

Quality control of the raw sequences from all 36 gill samples was performed using FASTQC (Andrews, 2010) to assess the initial quality. Subsequent quality improvement involved filtering out adaptor sequences and low-quality reads using TRIMMOMATIC, applying a phred +33 quality threshold, ensuring a minimum base quality score of 25, and retaining reads with a minimum length of 50 bp for downstream analyses (Bolger et al., 2014). A de novo assembly was constructed using TRINITY v2.9.1 (Grabherr et al., 2011) from the 36 pairs of clean sequences. Abundance estimation was carried out by mapping the reads of the raw nine paired sequences to the de novo transcriptome using SALMON (Patro et al., 2017). The completeness of the transcriptome assembly was evaluated with BUSCO v5.2.2 (Simão et al., 2015), utilizing the vertebrata_odb10 dataset (Creation date: 2021-02-19). An expression matrix was generated, filtering low-expression transcripts (minimum expression of 1.0) with TRINITY v2.9.1, and normalizing expression levels as transcripts per million transcripts (TPM). Likely protein-coding regions in transcripts were identified using TRANSDECODER v5.5.0 (Haas et al., 2013). The filtered transcriptome sequencing reads were aligned to protein databases, signal peptides, and transmembrane domains using DIAMOND v2.0.8 (Buchfink et al., 2015), SIGNALP 6.0 (Teufel et al., 2022), TMHMM v2.0 (Krogh et al., 2001), and HMMER v3.3.2 (Finn et al., 2011). Functional annotation of the de novo transcriptome was performed using TRINOTATE v3.2.2 (Grabherr et al., 2011).

### Differential Gene Expression Analysis

Prior to conducting differential gene expression analysis, clean reads from the six stations were aligned to the filtered transcriptome using SALMON to calculate the mapping rate. Mapping tables were merged according to stations and normalized using TMM (Robinson & Oshlack, 2010). Differential expression analysis was performed with three biological replicates for the gill tissues, comparing the reference station C to the rest of stations using the DESeq2 package (Love et al., 2014) with a fold change cutoff of >2 and FDR ≤0.05. This method identified significant gene expression differences between the gill tissues of all stations when compared the reference station, controlling for type I errors by incorporating FDR (Benjamini & Hochberg, 1995; J. J. Chen et al., 2010). Differentially expressed genes (DEGs) were visualized using a NMDS to explore the variation in the transcriptomic fingerprint of the different seasons, and the influence of the multiple stressors in their spatial variation. A GO pathway enrichment analysis was conducted over the differentially expressed genes DEGs using ShinyGo (Ge et al., 2020; version 0.8), applying FDR correction to account for multiple testing and control false positives (Khatri et al., 2012). The relevant pathways identified in the gene ontology and KEGG analysis were extracted to the database in supplementary material (S1). A canonical correspondence analysis CCA was performed to assess the relation of the anthropogenic stress variables and the magnitude of enrichment of the differentially expressed pathways. Assembly was executed on the high-performance computing system at the University of Cologne (CHEOPS), and the remaining analyses were performed using the Galaxy project platform (The Galaxy Community et al., 2022).

## Results

### Sequence quality control, de novo assembly, and annotation

The RNA sequencing conducted on gill tissue yielded an average of 33.48 (± 7.46 SD) millions of reads after quality control. The assembly with all sequences produced a raw transcriptome of 551.7 mb with an N50 of 2960 bp. The number of putative genes for the Trinity assembly was 265124, with a total transcript number of 416156. 3358 complete and fragmented BUSCOs were identified with an annotation completeness of 92.2%. Following the filtration of low-expression transcripts, 38.99% of the initial raw transcriptome was retained, resulting in a filtered transcriptome of 216 mb, encompassing 108328 putative genes with an N50 of 3092 bp. Notably, 2314 complete BUSCOs were retained, with an assembly completeness of 63.5%. Out of the filtered de novo transcriptome, 10000 transcripts were annotated using TRINOTATE, representing 61.62% of the total transcriptome.

### Differential gene expression analysis

When examining differential gene expression across the gradient stations and the reference station C, significant numbers of differentially expressed genes (DEGs) were identified for each station in both seasons. Overall, gene expression responses were stronger in summer, with a higher number of DEGs compared to the cold season. Station AID showed the strongest response in both seasons, with 1267 DEGs in the warm season and 770 DEGs in the cold season, likely due to higher stress levels at this station. In the warm season, station PHE had 622 DEGs, station A had 433 DEGs, station B had 585 DEGs, and station D had 971 DEGs. In the cold season, station PHE had 369 DEGs, station A had 653 DEGs, station B had 418 DEGs, and station D had 364 DEGs.

Non-metric Multidimensional Scaling (NMDS) analysis of the DEGs revealed significant changes in global gene transcription profiles induced by anthropogenic stress in *Cottus rhenanus* gills during both sampling campaigns. For both seasons, DEGs from biological replicates at different sites formed distinct groups in the NMDS plots (Figure 4 and Figure 5), confirming the quality and reliability of our sequencing. Notably, stations PHE and AID grouped together for both seasons, indicating a similar DEG signature. In the warm season (Figure 4), PHE and AID DEGs were influenced by shading, water level, dissolved oxygen concentration, and water temperature. In the cold season (Figure 5), PHE and AID DEGs were influenced by shading, water level, conductivity, and the concentrations of sulfate and chloride. This grouping pattern suggests that PHE and AID share a similar physiological response to high levels of anthropogenic stress.

**Figure 4.**
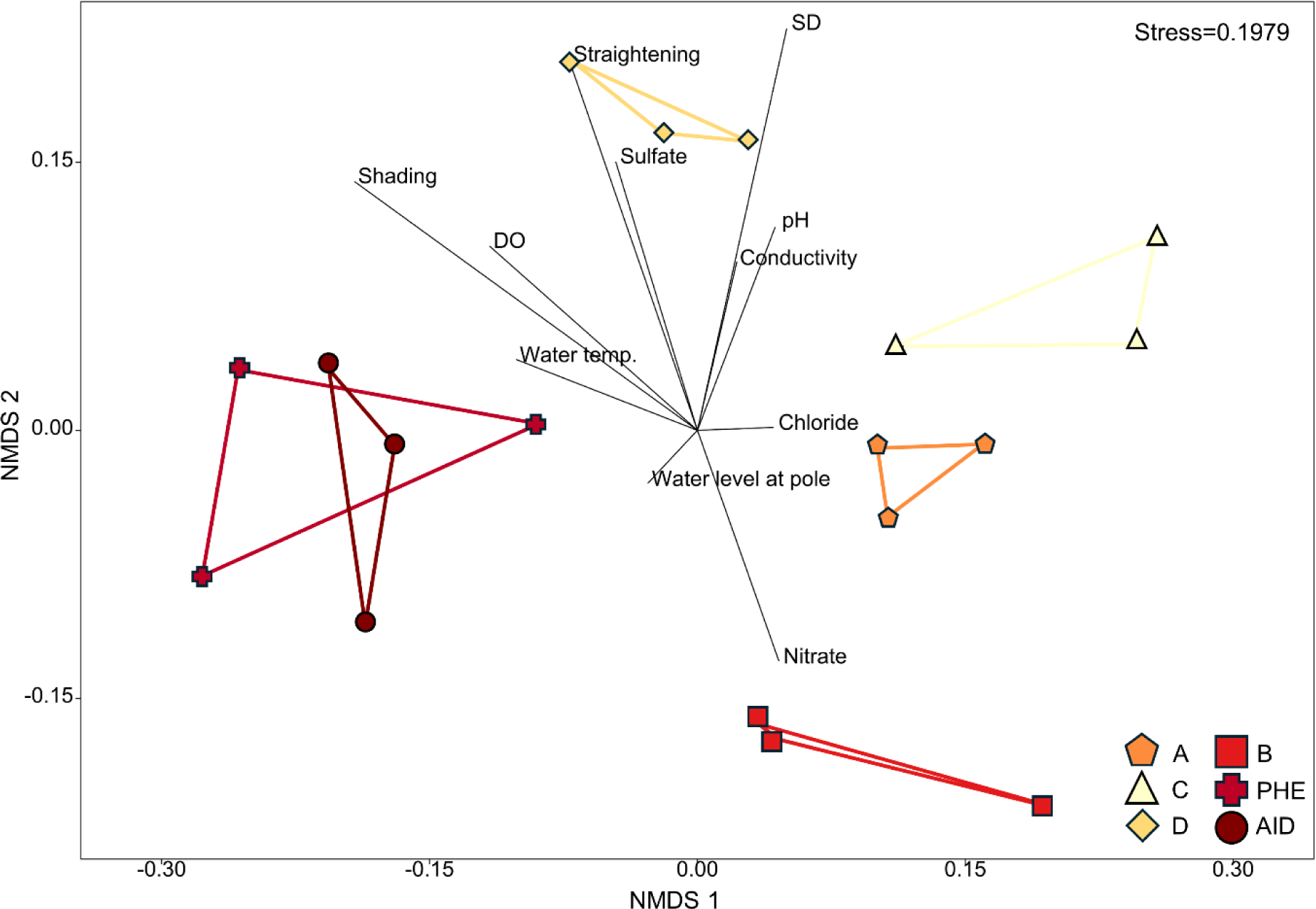
NMDS plot of DEGs and environmental variables in Emscher River stations (Warm season). Stress level is below the confidence threshold of 2.0 (0.1979). Biological replicates for each station form distinct groups. DEGs in Stations PHE and AID show similarity, influenced by shading, water level, dissolved oxygen, and water temperature. Station C samples group opposite the high stressor level stations PHE and AID.

**Figure 5.**
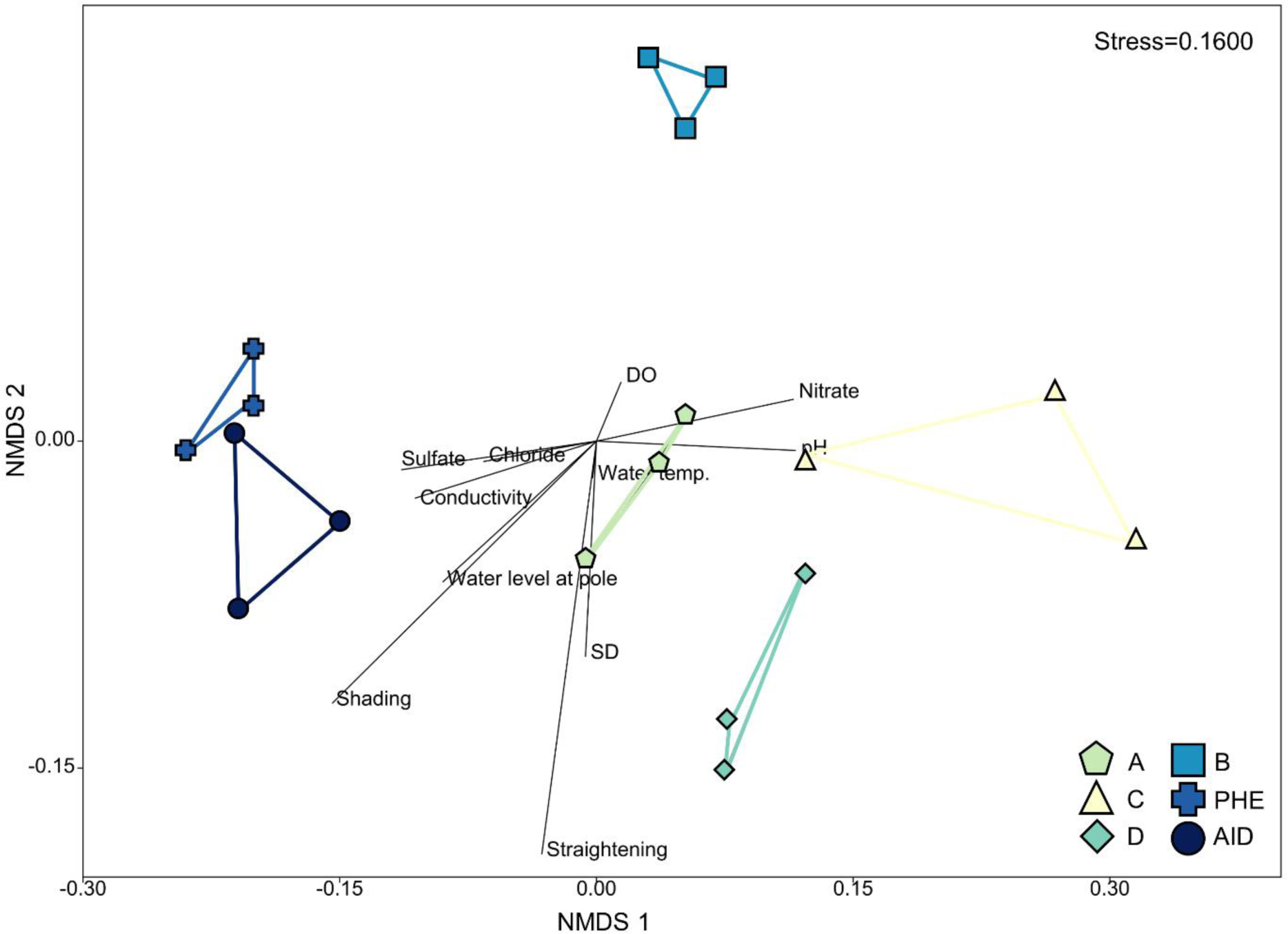
NMDS plot of DEGs and environmental variables in Emscher River stations (cold season). Stress level is below the confidence threshold of 2.0 (0.1600). Biological replicates for each station form distinct groups. DEGs in Stations PHE and AID show similarity, influenced by shading, water level, conductivity, sulfate, and chloride. Station C samples group opposite the high stressor level stations PHE and AID.

In both seasons, station C’s DEGs were on the opposite side of the NMDS plots from the PHE-AID group, reflecting the lower levels of anthropogenic stress at station C and validating its use as the reference station for differential gene expression analysis. Furthermore, stations A, B, and D were positioned at an intermediate distance between the PHE-AID group and station C. This organization can be explained by their intermediate levels of stress along the gradient between the reference conditions at station C and the highest anthropogenic stressors at stations PHE and AID. Notably, the stress values in both NMDS analyses were below the threshold of 2.0, allowing cautious interpretation of the relationships between DEGs and stressors. In summary, the differential gene expression patterns across the stations clearly reflect the varying levels of anthropogenic stress, with distinct grouping and stressor influences validated through NMDS analysis.

Overall, pathway enrichment was stronger during the warm period, with more DEGs enriched in both the KEGG and Gene Ontology databases across all samples. During the warm period, DEGs were significantly enriched (FDR p<0.05) for 2444 terms in the KEGG and Gene Ontology biological process, cellular component, and molecular component pathways. In the cold period, DEGs were significantly enriched (FDR p<0.05) for 1108 terms in the same pathways. Station AID had the most significantly enriched pathways in both seasons, with 809 in the warm period and 352 in the cold period. Conversely, station A had the fewest enriched pathways in the warm period (173), and station PHE had the fewest in the cold period (52) (Supplementary material S1).

The CCA explained 54.53% of the variation in enriched pathways across the different stations in relation to anthropogenic stressors during the warm season (CCA1: 28.73%, CCA2: 25.8%) (Figure 6). Enriched pathways for stations AID and PHE, the most stressed stations, were grouped together. This grouping was influenced by high water temperature, elevated chloride concentrations, high conductivity, and lower levels of pH, dissolved oxygen, and substrate diversity. Station D’s enriched pathways were influenced by higher nitrate and dissolved oxygen levels, and lower sulfate and conductivity levels. Stations A and B’s pathways were associated with higher pH, substrate diversity, sulfate, and conductivity. The CCA explained 61.32% of the variation in enriched pathways across the different stations in relation to anthropogenic stressors during the cold season (CCA1: 31.36%, CCA2: 29.96%) (Figure 7). Enriched pathways for stations AID, PHE, and B, the most stressed stations, were grouped together. This grouping was influenced by high water temperature, elevated chloride and sulfate concentrations, high conductivity, and lower levels of pH and nitrate. Stations D and A were characterized by higher levels of pH, substrate diversity, nitrate, and dissolved oxygen. In summary, CCA analysis revealed distinct patterns of pathway enrichment linked to specific anthropogenic stressors, varying between the warm and cold seasons. Stations AID and B were consistently influenced by variables related to anthropogenic stress in both seasons.

**Figure 6.**
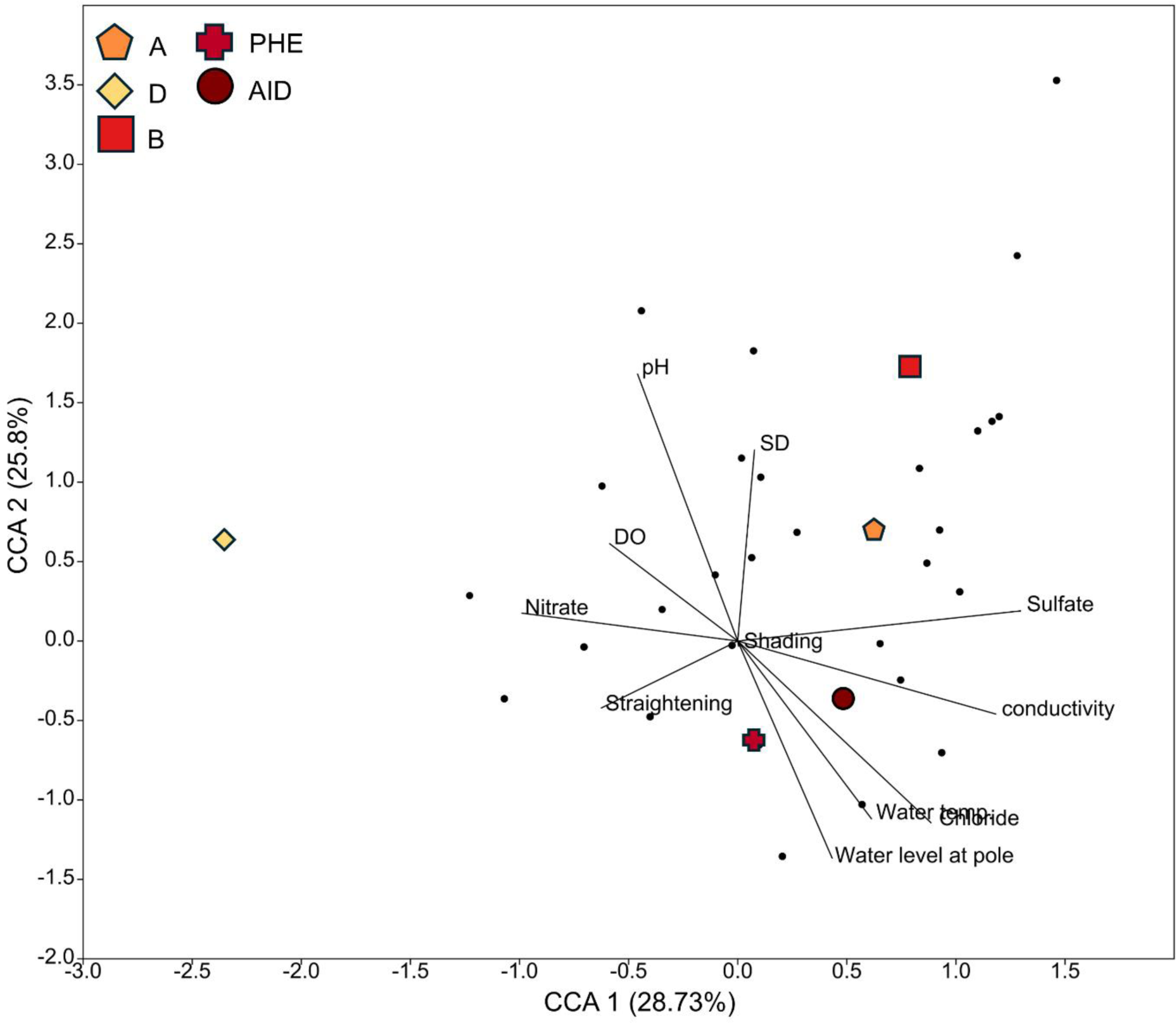
CCA plot of enriched pathways, and anthropogenic stressors for sampling stations during the warm season. The vectors represent the anthropogenic stressors. In colour and different geometrical shapes are the stations. The small black dots represent the gene ontology categories.

**Figure 7.**
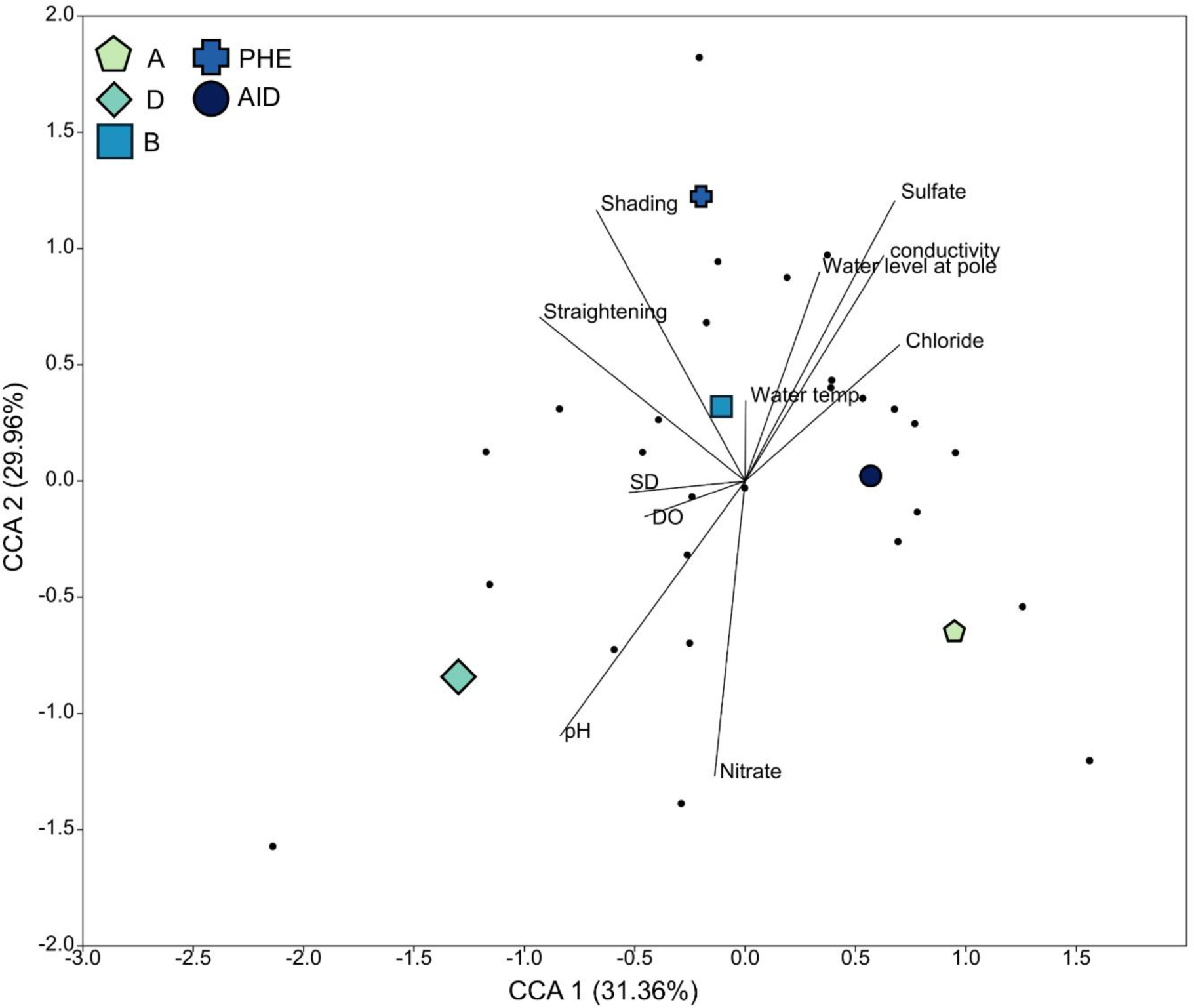
CCA analysis of KEGG, gene ontology categories, and anthropogenic stressors for sampling stations during the cold season. The vectors represent the anthropogenic stressors. In colour and different geometrical shapes are the stations. The small black dots represent the gene ontology categories.

Shared enriched pathways between stations AID and PHE encompass key parent GO groups such as Immune system process, regulation of transport, response to stress, and chemicals (Figure 8). Unique to station AID are pathways involving regulation of the MAPK cascade (GO:0043408, GO:0032872), regulation of ion transport (GO:0043270, GO:1901379) and circadian temperature homeostasis (GO:0060086), alongside distinct pathways related to immune system processes and stress responses (GO:0050776, GO:0002252). Conversely, exclusive pathways identified in station PHE include those related to immune system processes (GO:0006952), growth (GO:0040007), and reproduction (GO:0000003). Noteworthy, the hyperosmotic salinity response pathway (GO:0042538) and renal system development (GO:0072001) were enriched, with many genes relevant to transport and osmoregulation. Genes associated with regulation of transport include *ANO6*, *ANO9*, *KCNH1*, *KCNC2*, *WNK2*, *WNK4*, and *ABCA1*, reflecting roles in osmoregulation, ion transport, membrane dynamics, signalling, and cellular organization. In chemical response pathways, genes participate in responses to inorganic substances, nitrogen compounds, and metal ions. Within the response to stress category, pathways such as inflammatory response, defence response, and MAPK cascade are enriched. Finally, the immune system process category prominently features pathways related to lymphocyte and leukocyte functions, indicative of significant immune response gene involvement. This heatmap analysis highlights the distinct and overlapping genetic responses of stations AID and PHE to anthropogenic stress during the warm season, emphasizing the overexpression of immune-related pathways, osmoregulation and transport pathways, and overall metabolism related to growth and reproduction.

**Figure 8.**
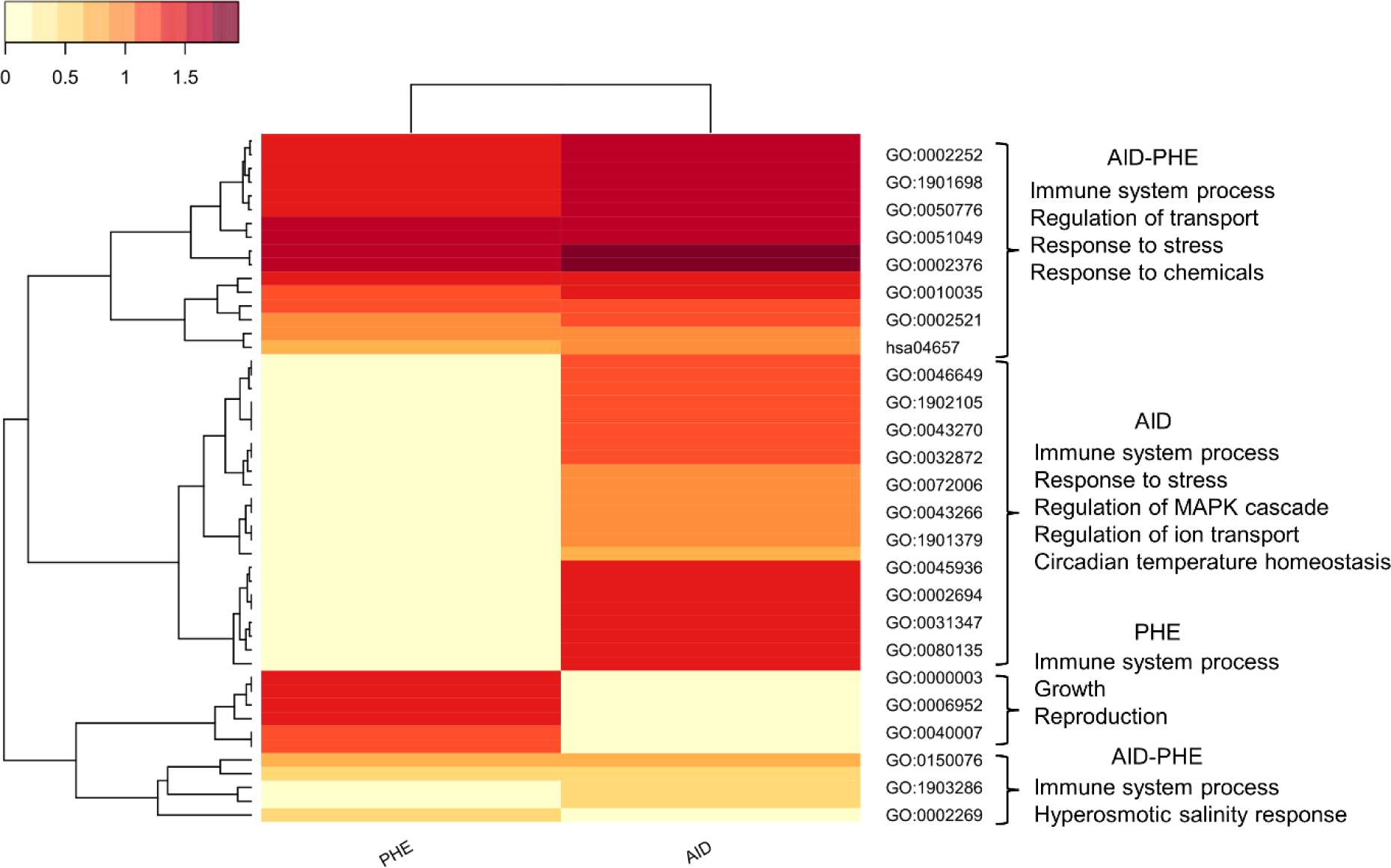
Heatmap of the number of genes for gene ontology and KEGG categories in high anthropogenic stress stations during the warm season. Stations AID and PHE are grouped based on shared gene ontology characterization and the number of genes per category. Brackets highlight the main categories found to be shared or unique to each station. The logfold number of genes has been transformed using Log10+1 for visualization.

Figure 9 presents a heatmap illustrating the distribution of genes across Gene Ontology (GO) and KEGG categories in high anthropogenic stress stations during the cold season, specifically stations AID, PHE, and B. The analysis reveals that the shared enriched pathways among these stations are primarily related to the parent GO category of developmental process. Station AID exhibits several unique pathways that reflect the influence of anthropogenic stressors, including response to stress, metabolic process, response to chemicals, and response to starvation. AID is also characterized by pathways related to the nervous system. Within these unique pathways, key genes associated with the response to oxidative stress pathway (GO:0006979) and overall response to stress (GO:0033554, GO:0006950) include *NFAT5*. The response to starvation pathway (GO:0042594, GO:0009267) was enriched in the station AID. For the response to chemicals category, genes belong to pathways related to oxidative stress (GO:0034599, GO:0000302) and organic compound metabolic processes (GO:0006807, GO:0071704). Shared pathways between all three stations fall under the parent category developmental process (GO:0032502), with many pathways related to nervous system development (GO:0010721, GO:0048699, GO:0007399) and tissue development (GO:0007275, GO:0009888). The metabolic process category includes genes in pathways related to primary metabolic processes (GO:0008152, GO:0044238), with key marker genes such as *ACADS* and ACADL. Additionally, the pathway respiratory gaseous exchange by the respiratory system (GO:0007585) is notably enriched in station B. This heatmap analysis underscores the distinct and overlapping genetic responses of stations AID, PHE, and B to anthropogenic stress during the cold season, highlighting specific pathways and genes involved metabolism and stress response mechanisms.

**Figure 9.**
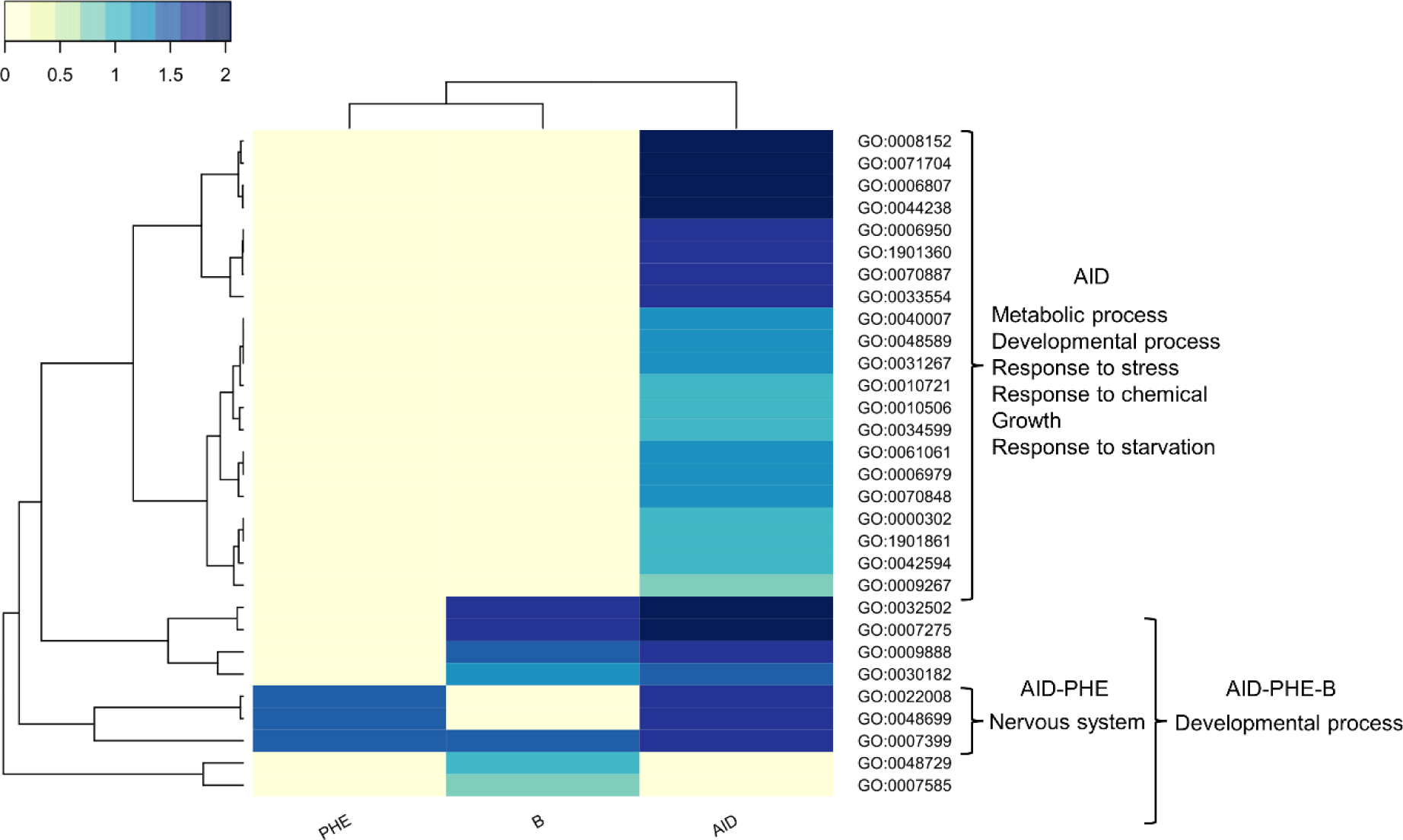
Heatmap of the number of genes for gene ontology and KEGG categories in high anthropogenic stress stations during the cold season. Stations AID, B and PHE are grouped based on shared gene ontology characterization and the number of genes per category. Brackets highlight the main categories found to be shared or unique to each station. The logfold number of genes has been transformed using Log10+1 for visualization.

The prevalence of immune response pathways in GO ontology for both PHE and AID prompted an in-depth analysis of enriched KEGG pathways using differentially expressed genes from both stations. This examination revealed pathways indicative of physiological stress, controlled by the expression of related genes. Notably, the MAPK (map04010) and IL-17 (hsa04657) signaling pathways were significantly influenced by environmental stress in the CCA analysis and included several DEGs. In the IL-17 signaling pathway, key DEGs identified were *IL1B* (Interleukin-1 beta, IL-1β), *TRAF6* (TNF Receptor Associated Factor 6), *NFKB1* (Nuclear Factor Kappa B Subunit 1), and *SOCS3* (Suppressor of Cytokine Signaling 3) (Figure 10). In the MAPK pathway, the significant genes included *IL1B* (IL-1β), *NFKB1* (NF-kappa-B p105/p50), *TRAF6, JUNB* (Transcription Factor Jun-B), *EGR1* (Early Growth Response Protein 1), ATF4 (Activating Transcription Factor 4) and *SMAD4* (Mothers Against Decapentaplegic Homolog 4) (Figure 11). Notably, *IL1B*, *TRAF6*, and *NFKB1* were shared between both pathways, highlighting their importance as biomarkers in the response to environmental stress.

**Figure 10.**
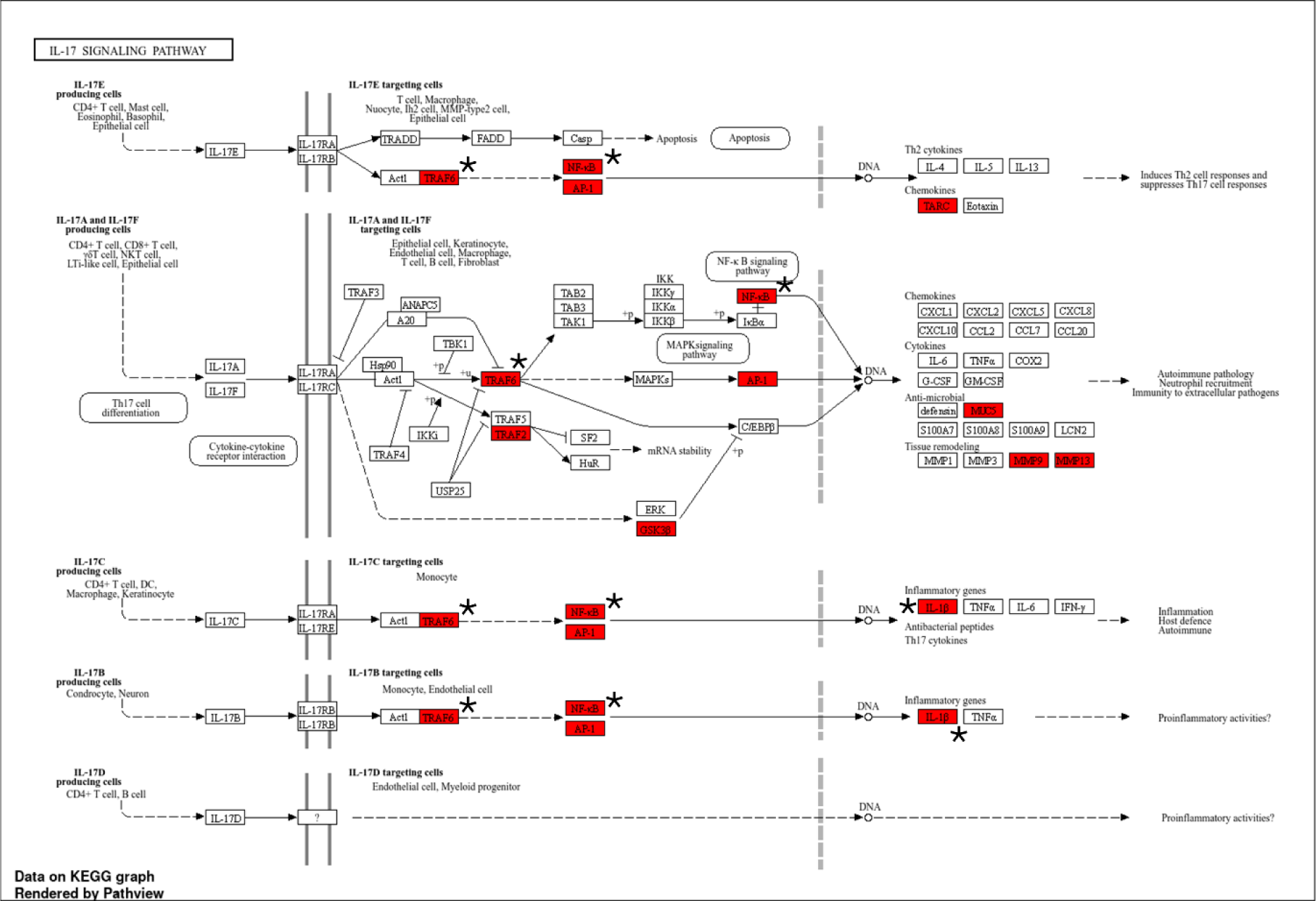
KEGG enriched pathway “IL-17 signalling pathway” for the gill tissue of *Cottus rhenanus*. Red highlighted boxes represent differentially expressed genes when comparing AID and PHE to reference station in summer. The genes marked with the symbol * are the ones shared between the IL-17 and MAPK signalling pathways.

**Figure 11.**
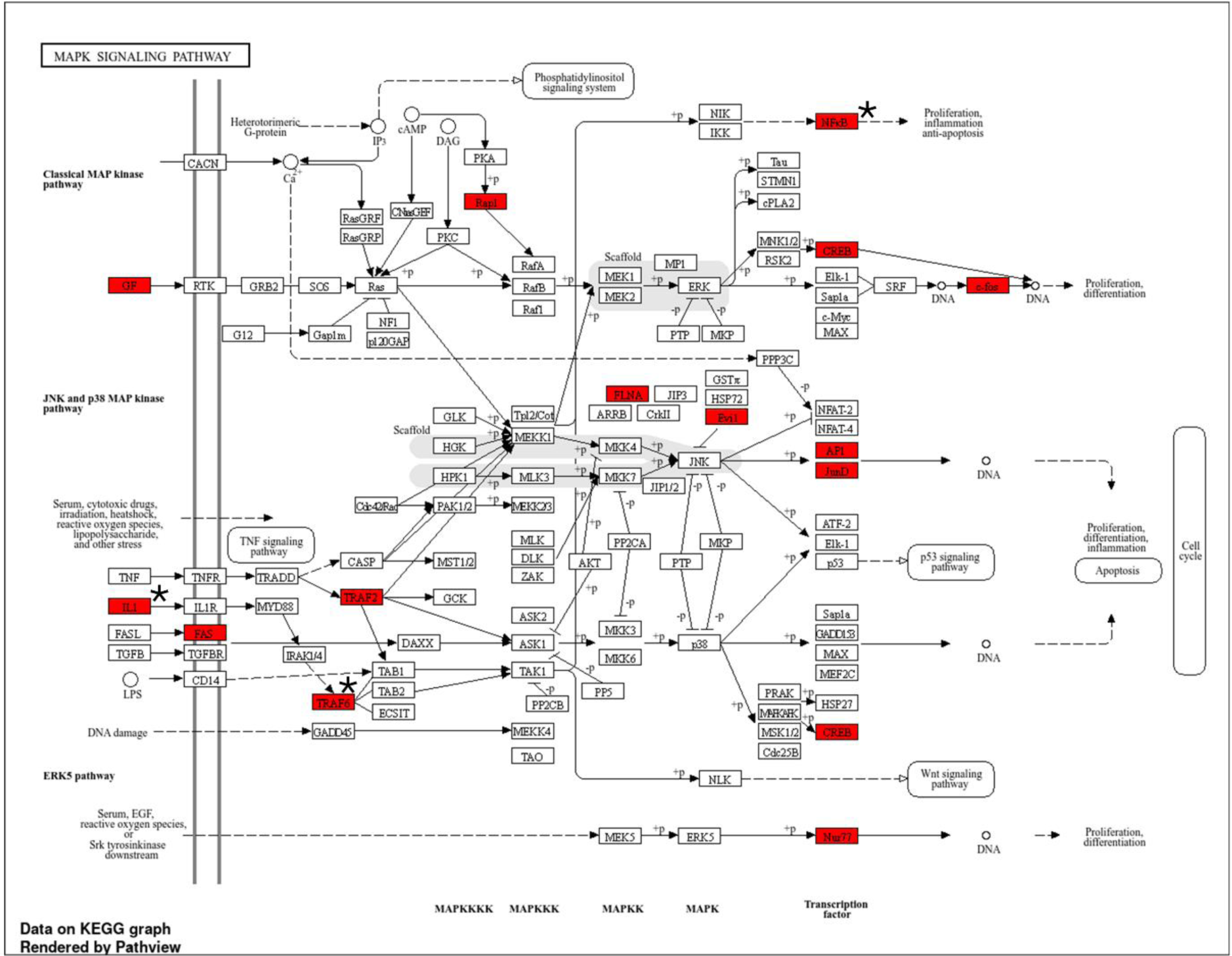
KEGG enriched pathway “MAPK signalling pathway” for the gill tissue of *Cottus rhenanus*. Red highlighted boxes represent differentially expressed genes when comparing AID and PHE to reference station in summer. The genes marked with the symbol * are the ones shared between the IL-17 and MAPK signalling pathways.

## Discussion

Our transcriptomic study offers a comprehensive analysis of *Cottus rhenanus* gills’ physiological responses to anthropogenic stress, emphasizing the study’s high data quality and key findings. High-quality RNA sequencing produced an average of 33.48 million reads per sample, resulting in a refined transcriptome assembly of 216 Mb with an N50 of 3092 bp, including 108,328 putative genes and 2314 complete BUSCOs, demonstrating 63.5% assembly completeness. Differential gene expression analysis revealed significant seasonal variations, particularly at Station AID during the warm season with 1267 DEGs, and NMDS analysis confirmed distinct grouping patterns of DEGs among stations, indicating consistent sequencing quality and reliability. The transcriptomic approach facilitates the inference of physiological responses in wild fish to a gradient of multiple stressors in an urban river. The study reveals that these stressors collectively impact the differential expression of genes associated with metabolism, transport, oxidative stress, and chemical response pathways, leading to an enhanced immune response. This is evidenced by a significant number of DEGs related to immune function and the enrichment of genes in the IL-17 and MAPK pathways at stations with higher anthropogenic stress during the warm season. Given the complexity of the enriched metabolic pathways and their interactions, the discussion focuses on pathways related to the main identified anthropogenic stressors, metabolism, osmoregulation and transport, and the overall immune response. The discussion concludes by highlighting the broader implications of the findings for research on multiple stressors and animal conservation.

## Metabolism

In this study the gill tissue of *Cottus rhenanus* from the most stressed stations during both cold and warm seasons, revealed significant enrichment of pathways related to metabolic stress, reproduction, and growth. Several recent studies assessing the transcriptomic response of fish to different stressors have found functional responses in pathways broadly categorized under growth and reproduction, consistent with our findings (Beemelmanns et al., 2021; Du et al., 2014; K. Zhou et al., 2020). Notably, during the warm season, there was a marked increase in differentially expressed genes (DEGs) involved in cellular metabolic processes. In the cold season, the response to starvation pathway was prominently enriched in stressed stations. The physiological responses observed align with previous studies highlighting strong metabolic reactions to similar stressors as it is discussed below.

One key stressor identified was salinity, particularly at stations with higher chloride concentrations and conductivities, where stress-related pathways were significantly enriched. Osmoregulatory processes are energy-intensive, involving lipid, glucose, and carbohydrate metabolism, as extensively reviewed by (Tseng & Hwang, 2008). Similar responses have been noted in other species; for instance, Milkfish (*Chanos chanos*) exhibit differential metabolic regulation in basal metabolism and oxidative phosphorylation under combined osmoregulatory and temperature stress (Hu et al., 2015). Additionally, hybrids of European minnow (Phoxinus sp.) show significant impacts on lipid metabolism under strong salinity stress (Escobar-Sierra et al., 2024). Temperature also influenced the stress transcriptomic response in both seasons, correlating with enriched pathways in lipid metabolism, as seen in juvenile turbot (*Scophthalmus maximus*) under temperature stress (T. Zhao et al., 2021). Lower dissolved oxygen levels in stressed stations further contributed to the enrichment of metabolic stress pathways, echoing findings in Yellow croaker (*Larimichthys polyactis*) (J. Wang et al., 2023).

Key biomarkers of metabolic stress were identified, including *ACADL* (Acyl-CoA Dehydrogenase Long Chain) and *ACADS* (Acyl-CoA Dehydrogenase Short Chain), which were differentially regulated in the AID station during the cold season. These genes, crucial in fatty acid metabolism, have shown differential expression under osmoregulatory stress in species such as *Eriocheir sinensis* and hybrid European minnows (Escobar-Sierra et al., 2024; Hui et al., 2014). The enrichment of the Cellular response to starvation pathway at the AID station during the cold season highlighted the stress endured. Notable genes within this pathway, such as *ATF4*, involved in glucose and lipid metabolism, were differentially expressed, consistent with findings in transgenic zebrafish and Japanese flounder under temperature stress (Han et al., 2023; Yeh et al., 2017). Additionally, *WDR24* and *NPRL3*, regulators of the mTORC1 pathway, were highlighted due to their roles in cell growth and nutrient signal integration. The activation of this pathway has been linked to metabolism imbalance under various stressors in species like *Oreochromis mossambicus*, European minnows, Chinese perch, and chinook salmon (Escobar-Sierra et al., 2024; Su et al., 2023; Tomalty et al., 2015; P. Wu et al., 2020). During the summer, *SOCS1* and *SOCS3* genes, biomarkers of metabolic stress regulating insulin signaling and glucose metabolism, were notably differentially expressed in the AID station, aligning with findings in Arctic charr and Japanese flounder under dietary stress (Deng et al., 2018; Jørgensen et al., 2013).

These findings emphasize the significant impact of multiple stressors on the metabolic pathways of *Cottus rhenanus*, with osmoregulatory and temperature stress playing critical roles.

### Hypoxia and oxidative stress

In both cold and warm seasons, stations experiencing stress exhibited lower dissolved oxygen concentrations, influencing the enrichment of specific stress pathways associated with chemical responses and stress. Detailed analysis revealed significant enrichment of pathways and differentially expressed genes (DEGs) linked to oxidative stress, indicative of a hypoxic physiological response. Hypoxia induces chronic and acute stress responses in fish, leading to molecular, behavioral, morphological, physiological, and immunological changes (Abdel-Tawwab et al., 2019). The fish gill, crucial for gas exchange, undergoes significant remodeling and leads the organismal response to varying dissolved oxygen levels (Mitrovic et al., 2009; C.-B. Wu et al., 2017). Under hypoxic conditions, fish experience reduced oxygen consumption and accelerated oxygen transport (Honda et al., 2019). While bioindicators in fish remain within normal ranges under normal dissolved oxygen concentrations, acute hypoxic stress increases reactive oxygen species (ROS) production (Sun et al., 2020). Fish have evolved an antioxidant defense system comprising enzymes and non-enzymatic molecules to regulate ROS homeostasis and adapt to hypoxia (Luo et al., 2021; Zhu et al., 2013). Failure of this system under hypoxic stress can exacerbate oxidative stress, tissue damage, apoptosis, or necrosis, contributing to the significant enrichment of oxidative stress-related pathways and DEGs observed in the gill tissue of *Cottus rhenanus* during both cold and warm seasons.

Transcriptomic studies investigating fish responses to hypoxia consistently identify the activation of oxidative stress-related genes and immune pathways. For example, in the silver carp gill, numerous DEGs indicate robust immune response and oxygen transport signaling under hypoxia (X. Li et al., 2022). Similar findings are reported in the yellow croaker liver and pearl gentian grouper, where pathways related to oxidative stress, apoptosis, and immunity are enriched (Liang et al., 2022; J. Wang et al., 2023). Notably, significant enrichment of the “Respiratory gaseous exchange by respiratory system” pathway at station B hints to the impact of dissolved oxygen stress. These results align with our findings, where alongside oxidative stress markers, we observed enrichment of immune responses indicated by IL-17 and MAPK pathway activation. The gill’s role as an immune-competent organ with extensive mucosal surfaces, known as gill-associated lymphoid tissue, further supports these observations (Koppang et al., 2015). Additionally, oxidative stress’s detrimental effect on fish immune and physiological functions is well-documented (Biller & Takahashi, 2018; Makrinos & Bowden, 2016; J. Wang et al., 2023).

Key candidate genes involved in oxidative stress and hypoxia were identified in the gill tissue of stressed stations across both seasons. Here we highlight some of these genes along with references to studies that have found them differentially expressed in response to hypoxia.

Examples include *G6PD* (Glucose-6-Phosphate Dehydrogenase), responsible for NADPH production crucial for ROS detoxification, and CCS (Copper Chaperone for Superoxide Dismutase), which activates *SOD1* to neutralize superoxide radicals (Lai et al., 2022). BMAL1, a circadian clock gene, and *CCN1* (Cyr61), involved in angiogenesis and cellular responses to hypoxia, were also differentially expressed (R.-X. Wu et al., 2023; H. Zhang et al., 2023). Additionally, *CDK1* and *CDK4*, regulators of cell cycle progression, showed enrichment in stressed stations during both seasons (Druker et al., 2021; He et al., 2017). *TRAF2* and *TRAF6*, mediators in TNF receptor signaling linked to oxidative stress and inflammation, exhibited high expression in stations stressed during the warm season (Jiang et al., 2023; G. Zhang et al., 2016).

In summary, the observed low dissolved oxygen levels in the stressed stations are leading to critical oxidative stress and hypoxia response pathways and biomarkers in *Cottus rhenanus* in the Boye River.

### Osmoregulation and transport

Freshwater salinity, driven by historical saltwater intrusion and urbanization, is a major stressor for *Cottus rhenanus* in the Emscher catchment. This study, along with previous research on three-spine stickleback in the same region, highlights that even low salinity levels can act as significant stressors. Even during the cold season, when osmoregulation and transport pathway enrichment was not paramount, key biomarkers of osmoregulation stress were recorded. These biomarkers, consistently found in various environments and different wild freshwater fish species under salinization stress, include *ANO6*, *ABCD1*, *ABCB6*, *SLC46A2*, and *NFAT5*. Anoctamins, such as *ANO6*, are involved in calcium-activated chloride channels and phospholipid scramblases, supporting cell volume regulation. These have been shown to be differentially expressed under salinity stress (Escobar-Sierra & Lampert, 2024; Hammer et al., 2015; Taugbøl et al., 2022). *ABCD1* and *ABCB6* are part of the ABC gene family, the largest group of transmembrane transporter proteins in the human genome. They use ATP to facilitate the transport of various molecules across cellular membranes, maintaining cell homeostasis (Bieczynski et al., 2021). These proteins have been reported in previous transcriptomic studies of fish under salinization stress (Escobar-Sierra & Lampert, 2024). *SLC46A2*, like other solute carriers in teleost fish, is involved in transmembrane transport, generating counter-ion fluxes to maintain ion homeostasis while avoiding excessive membrane tension (Saric & Freeman, 2021; Verri et al., 2012). This gene was also expressed during salinity stress in the Emscher fort he three spine stickelback (Escobar-Sierra & Lampert, 2024). *NFAT5* is crucial in the cellular response to hypertonic stress, regulating genes that help cells adapt to osmotic changes, a response previously reported under osmotic stress (Escobar-Sierra & Lampert, 2024; Pan et al., 2024).

During the warm season, stations PHE and AID exhibited greater impacts of salinization, with pathway enrichment influenced by higher electrical conductivities and chloride concentrations. The enrichment of ion transport pathways and key components of the transportome, particularly in gill tissues, aligns with consistent findings across various transcriptomic investigations that are discussed extensively in (Escobar-Sierra & Lampert, 2024) and (Escobar-Sierra et al., 2024). At PHE and AID, transport-related pathways were dominant, and pathways related to renal activity and hyperosmotic salinity stress were activated. These stations showed a physiological response similar to fish exposed to strong anthropogenic salinity pulses, including enriched osmoregulatory, immunological, and metabolism-related pathways (Escobar-Sierra et al., 2024). Key biomarkers such as *ABCA1*, *AQP3*, *KCNC2*, *WNK2*, *WNK4*, *ANO6*, *ANO9*, and *KCNH* were enriched, indicating the activation of osmoregulatory and ion transport systems. WNKs (With-No-Lysine [K] kinases) are involved in ion transport, blood pressure regulation, and sodium-coupled chloride cotransport, playing a crucial role in regulating cell volume in response to osmotic stress (Kahle et al., 2010). KCNCs, like other voltage-gated Ca2+ channels, regulate ion and water transport, as well as Ca2+ signaling and entry in animal epithelia (J. Zheng & Trudeau, 2015). *AQP3*, part of the aquaporin gene family, facilitates water and solute transport in osmotic gradients and is differentially expressed in fish gill, kidney, and gut tissue in response to salinity stress across various species (Cutler & Cramb, 2002; Escobar-Sierra & Lampert, 2024; Giffard-Mena et al., 2007). *RAB21*, *RAB25*, and *RAB3A* are rab GTPases involved in vesicle transport and exocytosis. These small GTPases, including Ras and Rho, regulate ion homeostasis through direct interaction with ion channels, with their expression under salinity stress previously recorded in fish (Escobar-Sierra & Lampert, 2024; Pochynyuk et al., 2007). Notably, anoctamins *ANO6* and *ANO9*, and the ABC gene family member *ABCA1*, were also differentially expressed in the stressed stations during the cold season, underscoring their importance as candidate genes for assessing osmoregulation stress responses.

These findings reveal the critical role of the transportome in the adaptive response of freshwater fish to salinity changes, even at sublethal chloride concentrations. The study reveals the need for effective freshwater salinization control measures to conserve aquatic fauna in European streams. Salinization must be considered in any restoration efforts in the Emscher catchment before reintroducing endangered species like *Cottus rhenanus*. This pattern is likely applicable to other urban streams facing similar anthropogenic pressures. In summary, salinity stress significantly impacts the physiological responses of freshwater fish in the Emscher catchment, with key biomarkers indicating activation of osmoregulatory mechanisms regardless of seasonal variations. This highlights the urgent need for addressing freshwater salinization in conservation and restoration efforts.

### Immune system

This study shows the significant effects of various stressors, such as salinity, water temperature, and oxidative stress, on the physiological responses of *Cottus rhenanus* in the urbanized Emscher catchment. Notably, the IL-17 and MAPK pathways are prominently activated at the AID and PHE stations during the warm season. Historically, research has concentrated on immune responses in primary and secondary lymphoid organs like the head, kidney, and spleen (Ewart et al., 2005; Overturf and LaPatra, 2006; Jørgensen et al., 2011). However, mucosal organs such as the gills, crucial for gas exchange and electrolyte balance, also play vital roles in pathogen defence (Evans et al., 1999; Koppang et al., 2015). Gills contain extensive mucosal surfaces and form the gill-associated lymphoid tissue, comprising various immune cells such as lymphocytes, macrophages, and antibody-secreting cells (Salinas, 2015).

Abiotic factors, including temperature fluctuations, crowding, salinity changes, and oxygen levels, profoundly influence fish immune systems. Previous studies have demonstrated that salinity stress, in particular, alters immune system processes with a notable involvement of the IL-17 and MAPK pathways (Escobar-Sierra et al., 2024). Furthermore, the IL-17 pathway have been found to be consistently activated in response to various stressors, such as ammonia, pathogens, heat, hypoxia, and salinity (Bai et al., 2022; X. Chen et al., 2021; Liang et al., 2022; Zhong et al., 2023). While the MAPK pathway is also frequently implicated in responses to stressors such as hypoxia and temperature changes (Lai et al., 2022; Song & McDowell, 2021; Tian et al., 2019; Y. Zhou et al., 2020). The IL-17 pathway, crucial for inflammatory responses, is significantly enriched under stress conditions, underscoring its role in fish immune response. Similarly, the MAPK pathway, a conserved signal transduction route, responds to diverse extracellular stimuli and regulates processes such as proliferation, differentiation, development, stress response, survival, and apoptosis (Gehart et al., 2010). Our findings align with studies in teleost fish, showing MAPKs are involved in responses to stressors like heat, osmotic and oxidative stress, ionizing radiation, and inflammatory cytokines (W. Zheng et al., 2022).

The presence of several key genes relevant to the IL-17 and MAPK pathways at stressed stations during summer highlights the importance of these pathways in regulating stress responses to various abiotic and biotic stressors. Specifically, the shared genes between these pathways (*IL1B*, *TRAF6*, and *NFKB1*) illuminate their pivotal roles in mediating the body’s response to environmental and stress-related challenges. *NFKB1*, a central regulator in the IL-17 pathway, orchestrates the transcriptional response to IL-17 signaling and contributes to the production of inflammatory mediators. This gene’s interaction with the MAPK and JAK/STAT pathways is essential for fine-tuning immune responses and maintaining homeostasis. Previous studies have demonstrated that *NFKB1* is differentially expressed under various stress conditions in fish, highlighting its importance in managing immune responses and stress adaptation (Van Muilekom et al., 2023; Xie et al., 2024). *SOCS3*, another key gene in the IL-17 pathway, modulates inflammatory responses by providing negative feedback to cytokine signaling, preventing excessive inflammation and protecting tissues from damage (Alexander, 2002). *SOCS3*’s role in maintaining immune homeostasis and its differential expression under metabolic stress in fish species such as the arctic charr and Japanese flounder further supports its importance in stress management and tissue growth regulation (Deng et al., 2018; Jørgensen et al., 2013; S. Zhao et al., 2020).

IL-1β is a significant player in both the IL-17 and MAPK pathways. In the IL-17 pathway, IL-1β enhances Th17 cell differentiation and the expression of inflammatory genes, while in the MAPK pathway, it activates MAPKs to drive the production of inflammatory mediators. This dual role in regulating inflammation and stress responses highlight *IL-1β*’s importance in coordinating cellular reactions to environmental changes affection (Krasnov et al., 2020; Rebl et al., 2020; Syahputra et al., 2020). *TRAF6*, as an adaptor protein, is crucial for mediating inflammatory and immune responses in both pathways. In the IL-17 pathway, *TRAF6* facilitates NF-κB and MAPK activation, leading to pro-inflammatory gene expression. In the MAPK pathway, *TRAF6*’s role in MAPK activation influences inflammation, cell differentiation, and survival. The relevance of *TRAF6* in oxidative stress and its response to environmental factors such as hypoxia in fish emphasize its broad impact on stress responses and immune regulation (Z. Li et al., 2019; R. Wang et al., 2023).

In addition to the shared genes, the MAPK pathway components such as *JUNB*, *EGR1*, *ATF4*, and *SMAD4* further illustrate the pathway’s complexity. *JUNB* and *EGR1* are crucial for regulating gene expression in response to extracellular signals, impacting inflammation, proliferation, and stress responses (Tiwari et al., 2013). *ATF4*’s role in managing oxidative stress and protein homeostasis highlights its importance in cellular stress adaptation, while *SMAD4*’s cross-talk with MAPK pathways underscores its significance in regulating responses to external stimuli (R. Wang et al., 2023; G. Zhang et al., 2016).

In summary, the interplay between the IL-17 and MAPK pathways, particularly through the shared genes *IL1B*, *TRAF6*, and *NFKB1*, demonstrates a strong immune response to multiple stressors in more anthropogenic-influenced stations during the summer. These genes are important candidates for assessing stress in wild fish under multiple stressors, suggesting that targeting these pathways in stress response studies is crucial for monitoring the health of freshwater fish and their ecosystems. Future research should validate these findings across various species and ecosystems.

### Wider implications and future directions

#### Multiple stressors and physiological response

This study identifies the physiological responses of fish to stressors in an urbanized multiple-stressor scenario in the wild. By relating the enrichment of various physiological pathways to increasing anthropogenic stressors such as decreasing dissolved oxygen, freshwater salinization, and temperature stress, candidate genes for rapid assessment of wild fish health were identified. Utilizing transcriptomics technologies allowed a holistic assessment of fish responses to multiple stressors in real-life scenarios, providing insights into the interactive effects on their overall physiological responses. Our findings align with the “metabolic compensation” strategy described by (Sokolova, 2013), where fish respond to stress by reallocating energy to defence mechanisms and maintenance, often at the expense of growth and reproduction and that Petitjean et al. (2019) consider is the response to single stressors. This strategy helps maintain homeostasis and survival but may transiently compromise the immune system. However, the response we describe is more in line with the pattern described by (Tort, 2011) regarding the response of fish to chronic stress, common in anthropogenically impacted environments, results in sustained energy reallocation to stress responses, which often translated in an altered immune function.

Interestingly, our results challenge the “metabolic conservation” strategy proposed by Petitjean et al. (2019), which suggests that under multiple stressors, high energy demands for maintenance may lead to a metabolic shutdown. This discrepancy may arise because previous observations were based on controlled lab conditions with fewer and more acute stressors. Our approach offers empirical insights in realistic ecological conditions, addressing the call for more studies to understand the physiological endpoints of individuals exposed to multiple stressors.

Future research should aim to establish a unifying conceptual framework for the physiological response to multiple stressors, similar to the Asymmetric Response Concept (ARC) proposed for ecosystem responses (Vos et al., 2023). This framework should consider the allostatic load or intensity of the stressor, gene expression changes, energy expenditure for homeostasis, and recovery trajectories after exposure to stressors. Efforts by Gandar et al., (2017) and Petitjean et al. (2019) have laid the groundwork, but new studies using omics tools provide a more holistic assessment. With the rapid advancement and affordability of high-throughput technologies, the comprehensive understanding of multiple stressor physiological responses is within reach and should be prioritized as a research direction in the field.

#### Transcriptomics in conservation of freshwater fish

Assessing the overall health or physiological status of organisms has traditionally focused on model species under controlled conditions with few stressors. The advent of de novo transcriptomics now enables work with non-model species without published genomes. Advances in RNA-seq affordability and bioinformatics have expanded its applicability to environmental sciences. As many reviews have stated, we are closer to accurately been able to assess the physiological status of organisms using transcriptomics with enormous implications to conservation (Connon et al., 2018; Jeffries et al., 2021; Semeniuk et al., 2022; Torson et al., 2020). Significant steps have demonstrated its applicability in unraveling the pathways and mechanisms of stress in wild fish (Escobar-Sierra et al., 2024; Escobar-Sierra & Lampert, 2024; Jeffrey et al., 2023; Komoroske et al., 2016). Specifically, we discerned the effects and their weight on the physiological response of sublethal stressors in a real-life scenario with multiple stressors interacting. The candidate genes identified in this study could be used for rapid assessment of anthropogenic stress in *Cottus rhenanus*, guiding conservation strategies for this endangered species. Rapid screening tools using candidate gene expression from transcriptomic research are being actively explored for assessing wild fish health (Chapman et al., 2021; Jeffries et al., 2021). Furthermore, our study identifies freshwater salinization, dissolved oxygen, and water temperature as highly influential stressors affecting the physiological response and immune system of *Cottus rhenanus*. For conservation of current populations in the Emscher catchment or planning new reintroductions in restored sites, these stressors must be carefully managed.

This study highlights the value of transcriptomics in assessing the health of non-model species in real-world scenarios. Future research should focus on refining rapid screening tools using candidate genes and exploring the physiological responses to multiple stressors in various species and environments. This approach will enhance conservation strategies and improve the management of endangered species under multiple anthropogenic stressors.

## Conclusion

Our transcriptomic study provides a comprehensive analysis of the physiological responses of *Cottus rhenanus* to multiple anthropogenic stressors in an urban river system. By leveraging high-throughput RNA sequencing and robust bioinformatics analysis, we identified significant variations in gene expression related to metabolism, oxidative stress, osmoregulation, transport, and immune responses, particularly in response to seasonal changes. The findings reveal a complex interplay between various stressors, such as high temperatures, salinity, low dissolved oxygen, and chemical pollutants, that collectively impact fish health and stress resilience. This study underscores the importance of understanding the holistic physiological responses of fish to multiple, concurrent stressors in their natural environment. Our identification of key biomarkers and enriched pathways offers valuable insights for monitoring fish health and informing conservation strategies. Future research should aim to integrate these transcriptomic insights into broader ecological assessments and management practices to mitigate the impact of urbanization on aquatic ecosystems and support the conservation of endangered species like *Cottus rhenanus*.

## Acknowledgments

This paper benefited from the multiple discussions within the Collaborative Research Centre 1439 RESIST (Multilevel Response to Stressor Increase and Decrease in Stream Ecosystems; www.sfb-resist.de). Funding was provided by the Deutsche Forschungsgemeinschaft (DFG, German Research Foundation; CRC 1439/1, project number: 426547801). Special thanks to Gunnar Jacobs and Bernd Stemmer from the Emschergenossenschaft und Lippeverband: EGLV and the Bezirksregierung Arnsberg-Obere Fischereibehörde for the help with sampling and logistics. We thank the Regional Computing Center of the University of Cologne (RRZK) for providing support and computing time on the High high-performance computing (HPC) system CHEOPS. And finally, we thank the Cologne Center for Genomics (CCG), for their technical support on sample preparation and sequencing.

## Notes

### Competing Interest Statement

The authors have declared no competing interest.

